# Single-cell analysis of intestinal immune cells during helminth infection

**DOI:** 10.1101/746479

**Authors:** Laura Ferrer-Font, Palak Mehta, Phoebe Harmos, Alfonso Schmidt, Kylie M Price, Ian F Hermans, Franca Ronchese, Graham Le Gros, Johannes U Mayer

**Affiliations:** Malaghan Institute of Medical Research, Wellington, New Zealand; Maurice Wilkins Centre, Wellington, New Zealand

## Abstract

Single cell isolation from helminth infected intestines has been notoriously difficult, due to the strong anti-parasite type 2 immune responses that drive mucus production, tissue remodeling and immune cell infiltration. Through the systematic optimization of a standard intestinal digestion protocol, we were able to isolate millions of immune cells from heavily infected tissues. Using this protocol, we validated many hallmarks of anti-parasite immunity and analyzed immune cells from the lamina propria and granulomas during helminth development, as well as acute and chronic worm infection.

## Main text

Recent advances in single cell analysis have significantly increased our understanding of multiple diseases and cell types in different tissues^1,2^. However, many of these technologies require single cell suspensions as an input, which limits our assessment of difficult-to-process tissues^2–4^. One prominent example is the intestine, which is at the center of many research questions that focus on nutrient uptake^5^, host-microbiome interactions^6–8^, local and systemic immune tolerance^9–11^ and gastrointestinal diseases and infections^12–16^, but represents a challenging tissue to digest^17,18^.

The standard digestion procedure to isolate intestinal immune cells located in the small intestinal lamina propria consists of three steps^17–20^(Fig. 1a). First the intestinal segment of interest is collected, opened longitudinally to remove its luminal content, washed and cut into small pieces. These pieces then undergo several wash steps with EDTA containing wash buffers at 37 °C to remove the epithelial layer and make the lamina propria accessible for enzymatic digestion. Lastly, the tissue is enzymatically digested with collagenases (Collagenase VIII from *Clostridium histolyticum* being among the most popular) and later filtered to obtain a single cell suspension.

**Figure 1.**
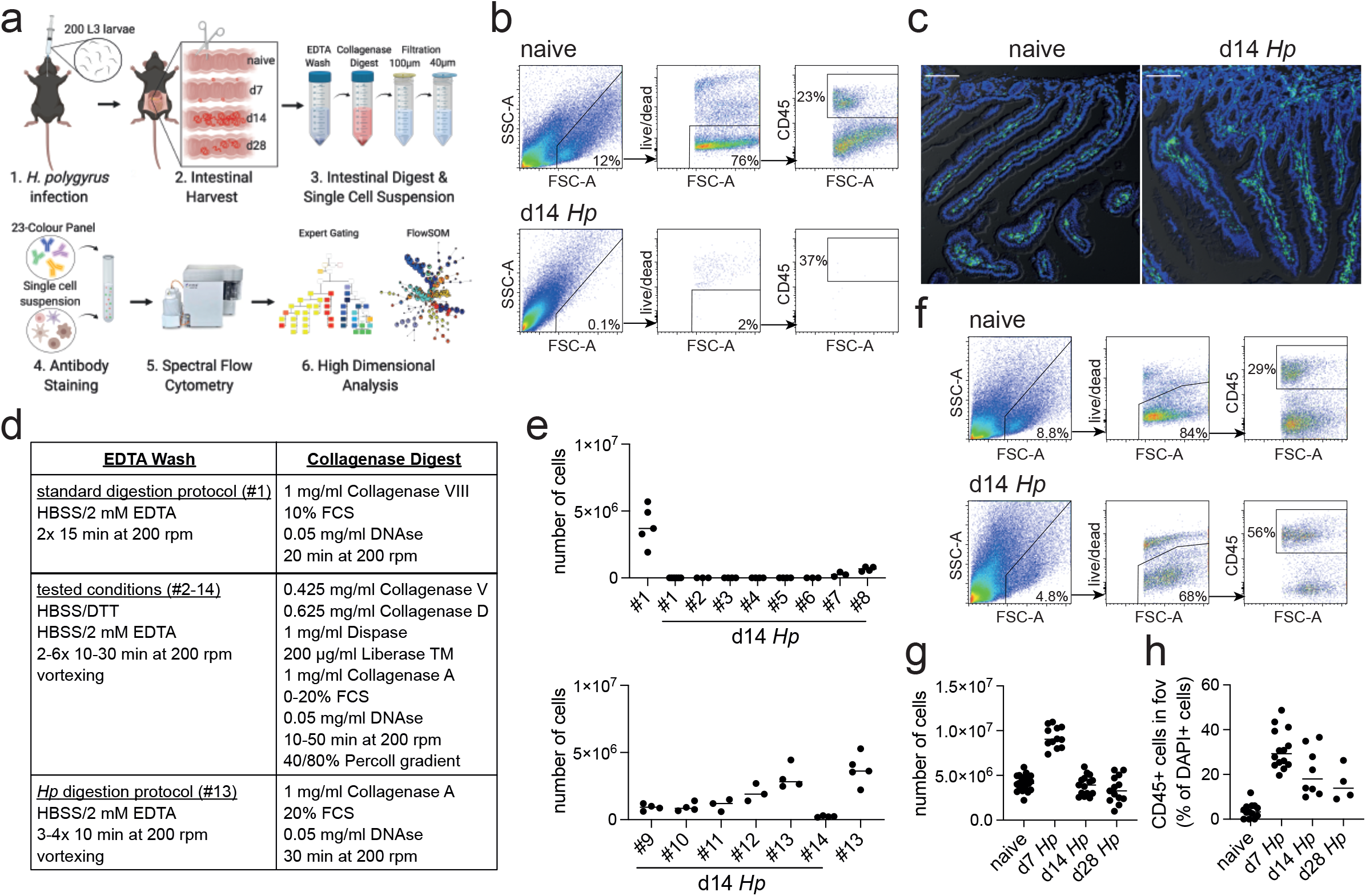
Optimization of a standard intestinal digestion protocol for heavily infected intestines. **a**, Schematic of a general intestinal digestion protocol (created with biorender.com). **b**, Failed digest of day 14 *H. polygyrus (Hp)* infected intestines using the standard digestion protocol. **c**, Intestinal cryosections stained with CD45-FITC (green) and DAPI (blue) from naïve and day 14 infected intestines. Scale bar=100μm. **d**, Summary of the different steps used for the systematic optimization of the standard digestion protocol. **e**, Number of live cells isolated from naïve or day 14 infected intestines for each of the tested conditions (for details see Supplementary Fig. 1,2) (n=3-5 samples per group, combined data from at least two independent experiments). **f**, Successful digest of day 14 infected intestines using the optimized *Hp* digestion protocol. **g**, Number of live cells isolated from naïve, day 7, day 14 and day 28 infected intestines using the optimized *Hp* digestion protocol (n>12 samples per group, combined data from at least three independent experiments). **h**, Quantification of CD45+ cells present in the field of view (fov) in cryosections from the same timepoints.

While this method results in a high cell yield from steady state intestines, the isolation of cells from severely infected segments remains challenging^18,19^(Fig. 1b). One of the most prominent examples are infections with intestinal helminths, which represent over 50% of all parasitic infections in human and life stock populations^21–24^. This has limited our analysis of antiparasite immune responses to imaging approaches, phenotypical observations in transgenic mouse strains or the assessment of secondary locations like the draining lymph nodes, blood or spleen^21,25,26^, which might only partially reflect local immunity. As helminth infections are strongly linked to chronic impairments that affect nutrition availability^27,28^; memory, cognition and physical development^29–31^; changes in the microbiota^32,33^and modulation of local and systemic immunity^26,34^, an optimized digestion protocol is needed to further investigate the infected intestinal tissue.

Difficulties with intestinal digests during helminth infection have been associated with a strong anti-parasite type 2 immune response that drives mucus production^35,36^, alters the epithelium^37,38^, induces immune cell infiltration^39^and causes tissue remodelling^40,41^(Fig. 1c). In order to investigate a model of both acute and chronic helminth infection, we infected C57BL/6 mice with *Heligmosomoides polygyrus*, a naturally occurring rodent parasite with an exclusive intestinal life cycle^42,43^. Infective L3 larvae penetrate the intestinal tissue of the duodenum within 24 hours of ingestion, undergo larval development in the muscularis externa and return to the lumen within 10 days post infection, where the adult worms mate and develop a chronic infection in C57BL/6 mice^43,44^. The peak of acute immunity is usually studied around day 14 post infection and we focused on this time point and the heavily infected duodenum^45,46^, to optimize our digestion protocol.

In order to develop a digestion protocol for heavily infected intestines we followed a systematic approach and optimized each step of the standard intestinal digestion protocol (Fig. 1d). First, we modified the EDTA wash steps to remove the increased amount of mucus but did not observe an improvement in cell yield (Fig. 1e and Supplementary Fig. 1; digestion protocols #1, 2, 6). This was followed by testing a variety of collagenases that have been reported for intestinal digests, as we hypothesized that the intestinal remodeling that occurred during helminth infection could negatively impact the digestion procedure. Indeed, we found that Collagenase A from *Clostridium histolyticum* (Fig. 1e and Supplementary Fig. 1; digestion protocols #7, 8), but not Collagenase VIII, Collagenase D, Dispase or Liberase TM (Fig. 1e and Supplementary Fig. 1; digestion protocols #3-5), showed an increase in cell yield when used in conjunction with the standard digestion protocol. To further optimize the protocol, we increased and modified the wash steps and observed a further increase in cell yield (Fig. 1e and Supplementary Fig. 2; digestion protocols #9-12). Importantly, strong vortexing after each wash step significantly improved the outcome of digestion (Fig. 1e and Supplementary Fig. 2; digestion protocol #13), suggesting that the epithelium is harder to remove in helminth infected tissues, likely due to its strong activation and remodelling^37,38^. Indeed, observations from Stat6ko mice confirmed that the physiological changes that impair the intestinal digest using the standard protocol, were all linked to type 2 immune responses, as intestines from infected Stat6ko mice could readily be digested (Supplementary Fig. 3). Several intestinal cell isolation protocols utilize a final gradient centrifugation step to further isolate immune cells^17,20^. However, in our hands this resulted in a dramatic drop in cell yield and was therefore omitted (Fig. 1e and Supplementary Fig. 2; digestion protocol #14). Our optimized lamina propria cell isolation protocol for *H. polygyrus* infected intestines thus included 3-4 10-minute 2mM EDTA wash steps (each followed by vigorous vortexing) and a 30-minute digest with 1mg/ml Collagenase A, 20% FCS and 0.05mg/ml DNase (see Supplementary Protocol 1 for step-by-step instructions).

**Figure 2.**
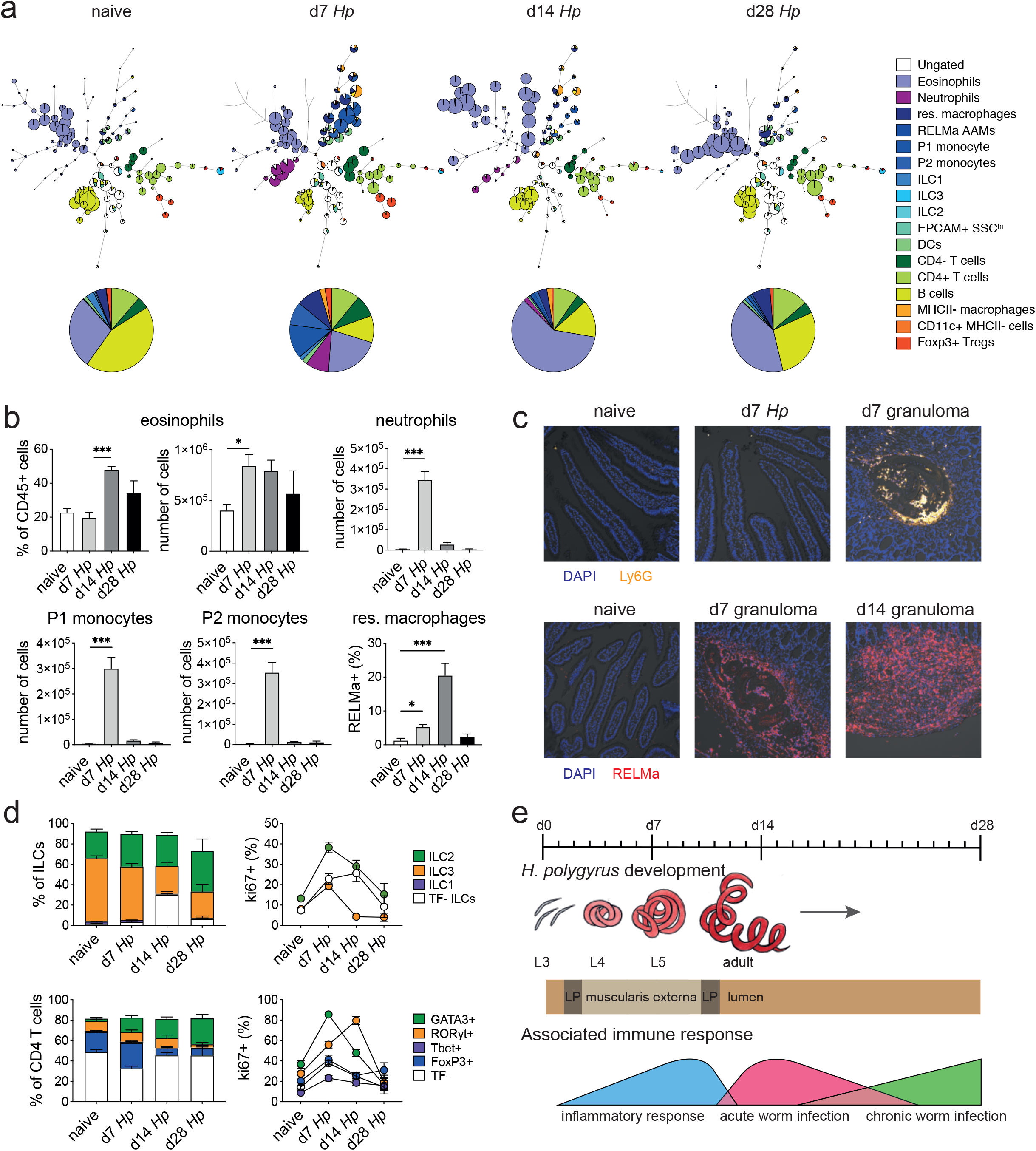
Spectral flow cytometric analysis of isolated intestinal immune cells during the course of *H. polygyrus* infection. **a**, FlowSOM (top) and manual (bottom) analysis of live CD45+ cells isolated from naïve, day 7, day 14 and day 28 infected intestines stained with 23 surface and intracellular antibodies and gated as described in Supplementary Fig. 6 (n=3-8 samples per group, combined data from two independent experiments). **b**, Quantification of infiltrating immune cells and RELMa+ macrophages during the course of *H. polygyrus* infection (mean±s.e.m.; Kruskal-Wallis followed by Dunn’s multiple comparisons test; *P≤0.05, ***P≤0.001). **c**, Representative images from intestinal cryosections stained with Ly6G-PECF594 (orange) and DAPI (blue) from naïve and day 7 infected intestines (top) or stained with RELMa-APC (red) and DAPI (blue) from naïve, day 7 and day 14 infected intestines (bottom) (representative of >10 sections from 3-5 mice per group and two independent experiments). **d**, Proportions of ILC and CD4 T cell populations and their expression of the proliferation marker ki67 during the course of infection. **e**, Schematic of *H. polygyrus* development, location and associated immune responses during the course of infection.

When we compared intestinal digests from naïve intestines using the standard or optimized cell isolation protocol, we observed highly comparable outcomes (Fig. 1e; digestion protocols #1 and 13). Both digestion protocols resulted in a cell yield of 3-6 million live cells per naïve duodenum with 70-80% viability and 20-30% frequency of CD45+ cells. To assess the effectiveness of our digestion protocol during the different stages of *H. polygyrus* infection, we harvested the duodenum from naïve C57BL/6 mice and at day 7, day 14 and day 28 post infection, which represented time points of larval development in the muscularis externa, acute as well as chronic adult worm infection, respectively. We observed that all time points could be successfully digested using our optimized digestion protocol and that duodenal digests from 14- and 28-days post infection yielded 3-6 million live cells per sample (Fig. 1g). We furthermore observed a consistent doubling of the cell count to 8-11 million live cells per duodenum at day 7 post infection and observed a similar trend when we quantified CD45+ cells in cryosections from these time points (Fig. 1g, h).

In order to understand these differences and validate that our protocol was suitable for subsequent single cell analysis and immunophenotyping, we characterized the isolated cells further using a 23-color spectral flow cytometry panel that incorporated many of the hallmark surface and intracellular markers for type 2 immune responses that have been associated with helminth infections^25,43^(see Supplementary Table 1 for details regarding markers, fluorophores, clones and staining concentrations used). As collagenase digests can negatively affect surface epitope integrity, we tested our optimized digestion protocol on splenocytes and compared digested to non-digested cells. While we observed a reduction in the MFIs of a few markers (namely Ly6G, MHCII, CD45 and CD127), all positive stained cell populations could be clearly identified (Supplementary Fig. 4). Isolated intestinal lamina propria cells also proved a challenge for intracellular staining, as different commercial intracellular staining kits significantly affected the cellular, but not debris, scatter profiles and varied in the resolution of intracellularly antibody staining (Supplementary Fig. 5). In our hands, the eBioscience FoxP3/Transcription Factor Staining Buffer Set yielded the best results and was used henceforth.

We used a combination of high-dimensional analysis tools and manual gating strategies to analyze the isolated lamina propria cells from the three main stages of *H. polygyrus* infection (Fig. 2a and Supplementary Fig. 6 and 7). In line with previous findings^39^, we observed a strong infiltration of immune cells such as neutrophils and monocytes at day 7 post infection, which we verified in cryosections and were primarily localized around the developing larvae explaining the increase in total cell number (Fig. 2a-c and Supplementary Fig. 8). At later time points this inflammatory response receded, which is likely linked to the worms exiting the intestinal tissue and inhabiting the lumen. Peak expression of RELMa in resident macrophages, which is a hallmark for their alternative activation and wound repair responses^47,48^, was observed at day 14 and was again localized within the granulomas (Fig. 2b,c and Supplementary Fig. 8). While type 2 innate lymphoid cells did not increase in frequency over time, ki67 expression increased, suggesting their proliferation and activation (Fig. 2b), as previously described^36,49^. GATA3+ Th2 cells, important drivers of type 2 immunity^43,50^, could be observed at day 7 post infection, increased in frequency over time and showed high ki67 expression (Fig. 2b). Interestingly, ki67 expression strongly decreased at day 28 post infection for all cell types analyzed (Fig. 2b and Supplementary Fig. 9), which could be linked to the strong immunomodulatory properties reported during chronic worm infection^25,51^(Fig. 2e).

Importantly, we were able to characterize intestinal innate and adaptive immune responses for all stages of *H. polygyrus* infection using our optimized lamina propria cell isolation protocol, which has previously not been possible. We were able to analyze immune cell infiltrations in the villi as well as the granulomas and could observe strong inflammatory responses during *H. polygyrus* development in the muscularis externa, followed by strong acute Th2 responses and a dampening of the immune response later on. Many of these changes were specific to the infected tissue and were not observed to the same extent in the draining lymph nodes (Supplementary Fig. 10 and 11), highlighting the potential of our protocol for future studies that utilize current single-cell analysis tools. We furthermore anticipate that this method will be applicable to study the immune responses to other intestinal helminths or type 2 immune conditions, such as *Nippostrongylus brasiliensis* infection or food allergy models, and after determining the correct collagenase compositions, could be also applied to human samples.

## Supporting information

Supplementary Protocol 1

## Supplementary Material

**Supplementary Protocol 1. Lamina propria cell isolation protocol for*H. polygyrus* infected intestines.**

**Supplementary Table 1. 23-color spectral flow cytometry panel for the analysis of intestinal immune cells during helminth infection.**

**Supplementary Figure 1. Modification of a standard intestinal digestion protocol to isolate single cells from heavily infected intestines.**

Dots plots of acquired events from day 14 H. polygyrus infected intestines using digestion protocols #1-8 (representative of 3-5 samples per group from at least two independent experiments). The gating shows cells of interest, viability and CD45 staining.

**Supplementary Figure 2. Further optimization of a single cell isolation protocol from heavily infected intestines based on Collagenase A digestion.**

Dots plots of acquired events from day 14 H. polygyrus infected intestines using digestion protocols #9-14 (representative of 3-5 samples per group from at least two independent experiments). The gating shows cells of interest, viability and CD45 staining.

**Supplementary Figure 3. Goblet cell hyperplasia and tissue remodeling during***H. polygyrus* **infection require Stat6ko signaling and impair cell isolation.**

**a**, H&E stained FFPE (Formalin fixed paraffin embedded) sections from naïve C57BL/6 and day 14 infected C57BL/6 and Stat6ko mice (representative of >10 sections from 3-5 mice per group and two independent experiments). **b**, Worm counts from day 14 infected C57BL/6 and Stat6ko mice (n=5 mice per group, representative for two independent experiments). **c**,**d**, Dots plots of acquired events and number of live cells isolated from naïve C57BL/6 and day 14 infected C57BL/6 and Stat6ko mice using the standard digestion protocol (representative of 3-5 samples per group from two independent experiments). The gating shows cells of interest, viability and CD45 staining.

**Supplementary Figure 4. Assessment of epitope integrity of digested and non-digested splenocytes.**

**a**, Comparison of Collagenase A digested and non-digested splenocytes stained for each surface marker used in our 23-color spectral flow cytometry panel. Gates and percentages of positively stained populations are shown (representative of two independent experiments). **b**, Comparison of MFIs for populations shown in **a**.

**Supplementary Figure 5. Comparison of different commercial intracellular staining kits.**

Dots plots of Collagenase A digested lamina propria cells from naïve C57BL/6 mice stained with Zombie NIR, CD45, CD4, ki67, FoxP3 and RORyt using four different commercial intracellular staining kits following the respective manufacturers’ instructions (representative of two independent experiments).

**Supplementary Figure 6. Gating strategy for small intestinal lamina propria cells.**

Dots plots of Collagenase A digested lamina propria cells from day 7 *H. polygyrus* infected C57BL/6 mice stained with our 23-color spectral flow cytometry panel. The gating strategy was used to identify innate and adaptive immune cells that have been associated with helminth infection and used for manual analysis of all stages of infection.

**Supplementary Figure 7. FlowSOM analysis of small intestinal lamina propria cells.**

**a**, FlowSOM analysis of live CD45+ cells isolated from naïve, day 7, day 14 and day 28 infected intestines stained with 23 surface and intracellular antibodies and gated as described in Supplementary Fig. 6 (n=3-8 samples per group, combined data from two independent experiments). **b**, Range of expression of each antibody for the combined FlowSOM analysis shown in **a**. Due to the uptake of Zombie NIR by eosinophils, only live CD45+ Siglec F-cells are shown.

**Supplementary Figure 8. Intestinal cryosections highlight infiltration of innate immune cells around the granulomas.**

Representative images from intestinal cryosections stained with CD45-AF488 (green), RELMa-APC (red), CD64-PE (white), Ly6G-PECF594 (orange) and DAPI (blue) from naïve, day 7, day 14 and day 28 infected intestines and their quantification per field of view (fov) (representative of >10 sections from 3-5 mice per group and two independent experiments).

**Supplementary Figure 9. Analysis of isolated intestinal immune cells during the course of***H. polygyrus* **infection.**

Analysis of live CD45+ cells (**a**), B cells (**b**), CD4 T cells (**c**), Dendritic cells (**d**) and CD64+ cells (**e**) isolated from naïve, day 7, day 14 and day 28 infected intestines as gated in Supplementary Fig. 6 (n=3-8 samples per group, combined data from two independent experiments; mean±s.e.m.; Kruskal-Wallis followed by Dunn’s multiple comparisons test; **P≤0.01).

**Supplementary Figure 10. Spectral flow cytometric analysis of mesenteric lymph nodes during the course of*H. polygyrus* nfection**.

**a**, FlowSOM (top) and manual (bottom) analysis of live CD45+ duodenum draining mesenteric lymph node cells from naïve, day 7, day 14 and day 28 infected mice stained with 23 surface and intracellular antibodies and gated as described in Supplementary Fig. 11 (n=2-9 samples per group, combined data from two independent experiments). Number and percentage of live cells (**b**), CD45+ cells (**c**), B cells (**d**), innate immune cells (**e,f**), ILCs (**g**), T cells (**h**) and CD4 T cells (**i**) are shown (mean±s.e.m.; Kruskal-Wallis followed by Dunn’s multiple comparisons test; *P≤0.05, **P≤0.01, ***P≤0.001).

**Supplementary Figure 11. Gating strategy for mesenteric lymph nodes.**

Dots plots of mesenteric lymph node cells from day 7 *H. polygyrus* infected C57BL/6 mice stained with our 23-color spectral flow cytometry panel. The gating strategy was used to identify innate and adaptive immune cells that have been associated with helminth infection and used for manual analysis of all stages of infection.

## Methods

### Ethics statement

All animal experiments were carried out at the Malaghan Institute of Medical Research and adhered to the regulations of the Ministry of Primary Industries New Zealand (license 24432) and were approved by the Animal Welfare Ethical Review Board of the Victoria University of Wellington.

### Mice

C57BL/6 (C57BL/6JOIaHsd) and C57BL/6.Stat6KO mice were bred at the Malaghan Institute of Medical Research, Wellington, New Zealand. Mice were housed under specific pathogen free conditions and age-matched female adult animals were used in each experiment.

### *Heligmosomoides polygyrus* infection

Mice were infected with 200 L3 larvae by oral gavage at 6-8 weeks of age and intestines and draining lymph nodes were harvested at the indicated time points.

### Cell isolation

Lamina propria cells were isolated from the first 8 cm of intestine according to the standard (Fig. 1e and Supplementary Fig. 1; digestion protocols #1)^19^, modified (Fig. 1e and Supplementary Fig. 1,2; digestion protocols #2-14) or optimized (see Supplementary Protocol 1, Fig. 1e and Supplementary Fig. 2; digestion protocol #13) protocols described in this manuscript. Individual duodenum draining mesenteric lymph nodes^20,52^were digested with 100μg/mL Liberase TL and 100μg/mL DNase I (both from Roche, Germany) for 30 minutes at 37 °C and passed through a 70μm cell strainer.

### Conventional and spectral flow cytometry

For conventional flow cytometry cells were resuspended in 0.5ml of 20μg/ml DNase containing FACS buffer, stained with DAPI to identify dead cells, filtered and analyzed using a BD LSRFortessa SORP™flow cytometer. For spectral flow cytometry, intestinal and lymph node samples were washed in 200μL FACS buffer and incubated with Zombie NIR Fixable Viability dye (Biolegend) for 15 minutes at room temperature. After washing, cells were incubated with Fc block (clone 2.4G2, affinity purified from hybridoma culture supernatant) for 10 minutes followed by the incubation of surface antibodies (see Supplementary Table 1) for 25 minutes at 4°C in the presence of 20μg/ml DNase and Brilliant Buffer Plus (BD Biosciences). Cells were fixed and permeabilization with the FoxP3/Transcription Factor Staining Buffer Set (eBioscience) according to manufacturer’s instructions and incubated with intracellular antibodies (see Supplementary Table 1) for 45 minutes at 4°C. Cells were then resuspended in FACS buffer, filtered, and analyzed on a 3- laser Aurora spectral flow cytometer (Cytek Biosciences).

### Data analysis

FCS files were manually analyzed using FlowJo (v10.6, Tree Star) or evaluated with high-dimensional data analysis tools using Cytobank (v7.2, Cytobank Inc.). After compensation correction in FlowJo, single, live, CD45+ events were imported into Cytobank and transformed to arcsinh scales. FlowSOM analysis was performed on 1,200,000 concatenated lamina propria and 1,000,000 concatenated lymph node cells, with an equal distribution of samples. Different cluster analyses were performed and 121 clusters were identified as the most representative for both data sets.

### Imaging

For histological sections, 5μm FFPE (Formalin fixed paraffin embedded) sections were stained using a standard H&E protocol^53^and visualized using a BX51 microscope (Olympus) equipped with a 10X NA 0.3 objective. For confocal microscopy, samples were fixed in 4% PFA for 1 hours and rinsed in PBS. Samples were then snap-frozen in OCT compound (Tissue-Tek) using a Stand-Alone Gentle Jane Snap-freezing system (Leica Biosystems). Cryosections of 7μm were blocked with Fc Block (clone 2.4G2, affinity purified from hybridoma culture supernatant) for 1 hour and stained with CD45-FITC (clone 30-F11, Biolegend), CD64-PE (clone X54-5/7.1, Biolegend), Ly6G-PECF594 (clone 1A8, Biolegend) or RELMa-APC (clone DSBRELM, eBioscience) for 1 hour. For nuclear staining, sections were incubated with DAPI (2 μg/ml) for 10 minutes. Images were taken with an inverted IX 83 microscope equipped with a FV1200 confocal head (Olympus) using a 20X, N.A 0,75 objective. Images were acquired using the FV10-ASW software (v4.2b, Olympus) and analyzed with ImageJ^54^(v1.52n) or CellProfiler^55^(v3.1.8). Image quantification analysis was based in the spatial co-expression of immune cell markers and DAPI-positive nuclei.

### Statistical analysis

Experimental group sizes ranging from 3 to 5 animals were chosen to ensure that a two-fold difference between means could be detected with a power of at least 80%. Prism 6 Software (GraphPad) was used to calculate the s.e.m. and statistical differences between groups and samples were calculated using Kruskal-Wallis tests followed by the appropriate multiple comparisons, with P≤0.05 being considered as significant.

### Source Data

Lamina propria and lymph node data sets used for manual and FlowSOM analysis can be downloaded from flowrepository (http://flowrepository.org/id/FR-FCM-Z28B). All other datasets are available upon request.

## Contributions

LFF, PM, PH, AS and JUM performed experiments and analyzed the data. KMP, IFH, FR and GLG offered support, provided reagents and helped direct the study. JUM conceived and directed the project and wrote the paper with input and feedback from all authors.

## Acknowledgments

We would like thank the staff of the Biomedical Research Unit and the Hugh Green Cytometry Core of the Malaghan Institute of Medical Research for their assistance.

This work was supported by an ‘In Aid of Research’ grant from the Research For Life foundation awarded to JUM and by grants from the Hugh Green Foundation, Maurice Wilkins Centre and the Health Research Council awarded to KMP, IFH, FR and GLG.

## References

1. Svensson, V., Vento-Tormo, R. & Teichmann, S. A. Exponential scaling of single-cell RNA-seq in the past decade. Nat Protoc 13,599–604 (2018).

2. Hwang, B., Lee, J. H. & Bang, D. Single-cell RNA sequencing technologies and bioinformatics pipelines. Exp Mol Med 50,96–14 (2018).

3. Nguyen, Q. H., Pervolarakis, N., Nee, K. & Kessenbrock, K. Experimental Considerations for Single-Cell RNA Sequencing Approaches. Front Cell Dev Biol 6,108 (2018).

4. Chen, X., Teichmann, S. A. & Meyer, K. B. From Tissues to Cell Types and Back: Single-Cell Gene Expression Analysis of Tissue Architecture. Annu Rev Biomed Data Sci 1,29–51 (2018).

5. Kiela, P. R. & Ghishan, F. K. Physiology of Intestinal Absorption and Secretion. Best Pract Res Clin Gastroenterol 30,145–159 (2016).

6. Belkaid, Y. & Hand, T. W. Role of the microbiota in immunity and inflammation. Cell 157,(2014).

7. Tilg, H., Zmora, N., Adolph, T. E. & Elinav, E. The intestinal microbiota fuelling metabolic inflammation. Nat Rev Immunol 336,973–15 (2019).

8. Sekirov, I., Russell, S. L., Antunes, L. C. M. & Finlay, B. B. Gut microbiota in health and disease. Physiol Rev 90,859–904 (2010).

9. Worbs, T. et al. Oral tolerance originates in the intestinal immune system and relies on antigen carriage by dendritic cells. J Exp Med 203,519–527 (2006).

10. Harrison, O. J. & Powrie, F. M. Regulatory T cells and immune tolerance in the intestine. Cold Spring Harb Perspect Biol 5,a018341–a018341 (2013).

11. Whibley, N., Tucci, A. & Powrie, F. Regulatory T cell adaptation in the intestine and skin. Nat Immunol 20,386–396 (2019).

12. Mowat, A. M. & Agace, W. W. Regional specialization within the intestinal immune system. Nat Rev Immunol 14,667–685 (2014).

13. Sell, J. & Dolan, B. Common Gastrointestinal Infections. Prim Care 45,519–532 (2018).

14. Fletcher, S. M., McLaws, M.-L. & Ellis, J. T. Prevalence of gastrointestinal pathogens in developed and developing countries: systematic review and meta-analysis. J Public Health Res 2,42–53 (2013).

15. Hendrickson, B. A., Gokhale, R. & Cho, J. H. Clinical aspects and pathophysiology of inflammatory bowel disease. Clin Microbiol Rev 15,79–94 (2002).

16. Saleh, M. & Elson, C. O. Experimental inflammatory bowel disease: insights into the host-microbiota dialog. Immunity 34,293–302 (2011).

17. Weigmann, B. et al. Isolation and subsequent analysis of murine lamina propria mononuclear cells from colonic tissue. Nat Protoc 2,2307–2311 (2007).

18. Reißig, S., Hackenbruch, C. & Hövelmeyer, N. Isolation of T cells from the gut. Methods Mol Biol 1193,21–25 (2014).

19. Scott, C. L., Wright, P. B., Milling, S. W. F. & Mowat, A. M. Isolation and Identification of Conventional Dendritic Cell Subsets from the Intestine of Mice and Men. Methods Mol Biol 1423,101–118 (2016).

20. Esterházy, D. et al. Compartmentalized gut lymph node drainage dictates adaptive immune responses. Nature 535,75 (2019).

21. McSorley, H. J. & Maizels, R. M. Helminth infections and host immune regulation. Clin Microbiol Rev 25,(2012).

22. Hotez, P. J. et al. Helminth infections: the great neglected tropical diseases. J Clin Invest 118,1311–1321 (2008).

23. Jourdan, P. M., Lamberton, P. H. L., Fenwick, A. & Addiss, D. G. Soil-transmitted helminth infections. Lancet 391,252–265 (2018).

24. Jackson, J. A., Friberg, I. M., Little, S. & Bradley, J. E. Review series on helminths, immune modulation and the hygiene hypothesis: immunity against helminths and immunological phenomena in modern human populations: coevolutionary legacies? Immunology 126,18–27 (2009).

25. Maizels, R. M. & McSorley, H. J. Regulation of the host immune system by helminth parasites. J Allergy Clin Immunol 138,666–675 (2016).

26. Mishra, P. K., Palma, M., Bleich, D., Loke, P. & Gause, W. C. Systemic impact of intestinal helminth infections. Mucosal Immunol 7,753–762 (2014).

27. Koski, K. G. & Scott, M. E. Gastrointestinal nematodes, nutrition and immunity: breaking the negative spiral. Annu Rev Nut. 21,297–321 (2001).

28. Crompton, D. W. T. & Nesheim, M. C. Nutritional impact of intestinal helminthiasis during the human life cycle. Annu Rev Nutr 22,35–59 (2002).

29. Ezeamama, A. E. et al. Helminth infection and cognitive impairment among Filipino children. Am J Trop Med Hyg 72,540–548 (2005).

30. Pabalan, N. et al. Soil-transmitted helminth infection, loss of education and cognitive impairment in school-aged children: A systematic review and meta-analysis. PLoS Negl Trop Dis 12,e0005523 (2018).

31. Nokes, C., Grantham-McGregor, S. M., Sawyer, A. W., Cooper, E. S. & Bundy, D. A. Parasitic helminth infection and cognitive function in school children. Proc Biol Sci 247,77–81 (1992).

32. Gause, W. C. & Maizels, R. M. Macrobiota - helminths as active participants and partners of the microbiota in host intestinal homeostasis. Curr Opin Microbiol 32,(2016).

33. Ramanan, D. et al. Helminth infection promotes colonization resistance via type 2 immunity. Science 352,608–612 (2016).

34. Maizels, R. M., Pearce, E. J., Artis, D., Yazdanbakhsh, M. & Wynn, T. A. Regulation of pathogenesis and immunity in helminth infections. J Exp Med 206,2059–2066 (2009).

35. Hashimoto, K. et al. Depleted intestinal goblet cells and severe pathological changes in SCID mice infected with Heligmosomoides polygyrus. Parasite Immunol 31,457–465 (2009).

36. Moltke, von, J., Ji, M., Liang, H.-E. & Locksley, R. M. Tuft-cell-derived IL-25 regulates an intestinal ILC2-epithelial response circuit. Nature 529,221–225 (2016).

37. Gerbe, F. et al. Intestinal epithelial tuft cells initiate type 2 mucosal immunity to helminth parasites. Nature 529,226–230 (2016).

38. Howitt, M. R. et al. Tuft cells, taste-chemosensory cells, orchestrate parasite type 2 immunity in the gut. Science 351,1329–1333 (2016).

39. Inclan-Rico, J. M. & Siracusa, M. C. First Responders: Innate Immunity to Helminths. Trends Parasitol 34,861–880 (2018).

40. Motran, C. C. et al. Helminth Infections: Recognition and Modulation of the Immune Response by Innate Immune Cells. Front Immunol 9,664 (2018).

41. Boyett, D. & Hsieh, M. H. Wormholes in host defense: how helminths manipulate host tissues to survive and reproduce. PLoS Pathog 10,e1004014 (2014).

42. Monroy, F. G. & Enriquez, F. J. Heligmosomoides polygyrus: A model for chronic gastrointestinal helminthiasis. Parasitol Today 8,49–54 (1992).

43. Reynolds, L. A., Filbey, K. J. & Maizels, R. M. Immunity to the model intestinal helminth parasite Heligmosomoides polygyrus. Semin Immunopathol 34,829–846 (2012).

44. Smith, K. A. et al. Low-level regulatory T-cell activity is essential for functional type-2 effector immunity to expel gastrointestinal helminths. Mucosal Immunol 9,428–443 (2016).

45. Filbey, K. J. et al. Innate and adaptive type 2 immune cell responses in genetically controlled resistance to intestinal helminth infection. Immunol Cell Biol 92,436–448 (2014).

46. Elliott, D. E. et al. Colonization with Heligmosomoides polygyrus suppresses mucosal IL-17 production. J Immunol 181,(2008).

47. Esser-von Bieren, J. et al. Antibodies Trap Tissue Migrating Helminth Larvae and Prevent Tissue Damage by Driving IL-4Rα-Independent Alternative Differentiation of Macrophages. PLoS Pathog 9,(2013).

48. Krljanac, B. et al. RELMα-expressing macrophages protect against fatal lung damage and reduce parasite burden during helminth infection. Sci Immunol 4,eaau3814 (2019).

49. Schneider, C. et al. A Metabolite-Triggered Tuft Cell-ILC2 Circuit Drives Small Intestinal Remodeling. Cell 174,271–284.e14 (2018).

50. Mohrs, K., Harris, D. P., Lund, F. E. & Mohrs, M. Systemic Dissemination and Persistence of Th2 and Type 2 Cells in Response to Infection with a Strictly Enteric Nematode Parasite. J Immunol 175,5306–5313 (2014).

51. Grainger, J. R. et al. Helminth secretions induce de novo T cell Foxp3 expression and regulatory function through the TGF-β pathway. J Exp Med 207,(2010).

52. Mayer, J. U. et al. Different populations of CD11b+ dendritic cells drive Th2 responses in the small intestine and colon. Nat Commun 8,15820 (2017).

53. Fischer, A. H., Jacobson, K. A., Rose, J. & Zeller, R. Hematoxylin and eosin staining of tissue and cell sections. CSH Protoc 2008,(2008).

54. Schindelin, J. et al. Fiji: an open-source platform for biological-image analysis. Nat Methods 9,676–682 (2012).

55. Lamprecht, M. R., Sabatini, D. M. & Carpenter, A. E. CellProfiler: free, versatile software for automated biological image analysis. BioTechniques 42,71–75 (2007).

