## Supplementary Protocol 1 for "Single-cell analysis of intestinal immune cells during helminth infection"

### **Lamina propria cell isolation protocol for *H. polygyrus* infected intestines**

according to *Ferrer-Font et. al.* 2019

Reagents

- RPMI (Gibco RPMI 1640 Medium)
- HBSS (Gibco Hank's Balanced Salt Solution)
- UltraPure 0.5M EDTA (Thermofisher #15575020)
- DNAse I (Sigma #10104159001)
- Collagenase A from *Clostridium histolyticum* (Roche #10103578001)
- Fetal Bovine Serum (Gibco)
- DPBS (Gibco)
- FACS buffer (PBS, 2% FBS, 2mM EDTA)
- EDTA wash buffer (HBSS, 2mM EDTA)
- Wash buffer (HBSS)
- Collection buffer (HBSS, 2% FBS)
- Digestion mix (RPMI, 20% FBS, 1mg/ml Collagenase A, 0.05mg/ml DNAse)
- Trypan Blue Solution (Gibco)

Materials

- Funnels
- 10x10cm Gauze (140µm mesh size)
- Tweezers to hold gauze in place
- 50ml falcon tubes
- 40µm cell strainers
- 100µm cell strainer
- 25 ml serological pipette
- Scissor/tweezers for collection
- Petri dish
- Haemocytometer

Equipment

- Shaking incubator
- Centrifuge

Step-by-step protocol

1. Warm bottles of RPMI, HBSS and HBSS/2mM EDTA to 37 °C.

2. Prepare 50ml falcon tubes with 10ml HBSS/2% FBS for each sample (keep on ice). Take petri dish, PBS and scissors/tweezers for collection of samples.
3. Sacrifice mouse, spray with EtOH and perform midline incision of the skin and muscle layer (see panel A in figure below).
4. Expose the intestines and excise the entire small intestine or a segment of interest and separate it from mesentery/fat tissue (see panels B-D).
5. Place intestinal segment on moist paper and remove the Peyer's patches.
6. Cut the segment longitudinally (see panel E).
7. Remove worms and intestinal content (see panels F&G).
8. Wash intestinal segment in PBS and cut into small pieces (~5mm long) (see panels H&I).
9. Collect pieces in 50ml falcon tube containing 10 ml HBSS/FBS, shake well and keep on ice (see panel J).
10. Repeat steps 3-8 for remaining samples.
11. Prepare 10ml fresh digestion mix for each sample containing 20% FBS, 1mg/ml Collagenase A and 0.05mg/ml DNAse in RPMI and warm up to 37° C.
12. Filter each sample through a 10x10cm gauze (140µm mesh size) placed on top of a funnel. Discard the flow-through (see panel K).
    *The same gauze can be reused in each wash step.*
13. Wash sample with 10ml of warm HBSS twice. Discard the flow-through (see panel l).
14. Remove the gauze from the funnel and collect sample in a 50ml falcon tube containing 10 ml HBSS/EDTA (see panels M-O).
    *It is advisable to process no more than 4-5 samples in parallel.*
15. Incubate samples for 10 minutes at 37° C and 200rpm in a shaking incubator (see panel P).
16. Vigorously vortex sample for 10 seconds after incubation.
    *A cloudy suspension should be observed.*
17. Repeat wash steps 10-14 two more times.
    *The HBSS/EDTA suspension should become less cloudy with each wash step. If significant amounts of debris are still observed after the last wash add a forth wash step.*
18. After the final EDTA wash step, filter each sample through a 10x10cm gauze and wash with 10ml of warm HBSS twice.
19. Remove the gauze from the funnel and collect sample in a 50ml falcon tube containing 10ml digestion mix.
20. Incubate samples for 30 minutes at 37 °C and 200rpm in a shaking incubator. Shake samples vigorously every 5 minutes.
    *The pieces will not be fully digested at this time point but have been optimised to yield the highest cell number with the least effect on viability and epitope integrity.*
21. Add 10ml of FACS buffer to each sample and keep on ice to stop the digestion.

22. Filter each sample through a 100 and 40µm cell strainer into a new 50ml falcon tube using a 25ml serological pipette and place on ice (see panels Q&R).
    *Do not simply pipette the sample on top of the strainer but force it through the mesh with pressure from the pipette controller.*
23. Centrifuge the single cell suspension at 600 x *g* for 6 min at 4 °C.
24. Discard the supernatant and resuspend in 1ml of 20µg/ml DNAse containing FACS buffer.
    *Use DNAse containing FACS buffer in all downstream procedures to reduce clumping.*
25. Count live cells in a 1:1 suspension of Trypan Blue using a haemocytometer.
26. Process desired number of cells for flow cytometry or other downstream applications.
    *Lamina propria digests contain a high percentage of debris, which will stain with viability dyes. Increase the concentration of viability dye and record samples using a high Scatter threshold to facilitate the identification of cells. If cells are to be sorted, pre-enrich the samples using positive selection beads. Avoid enrichment systems that are easily clogged.*

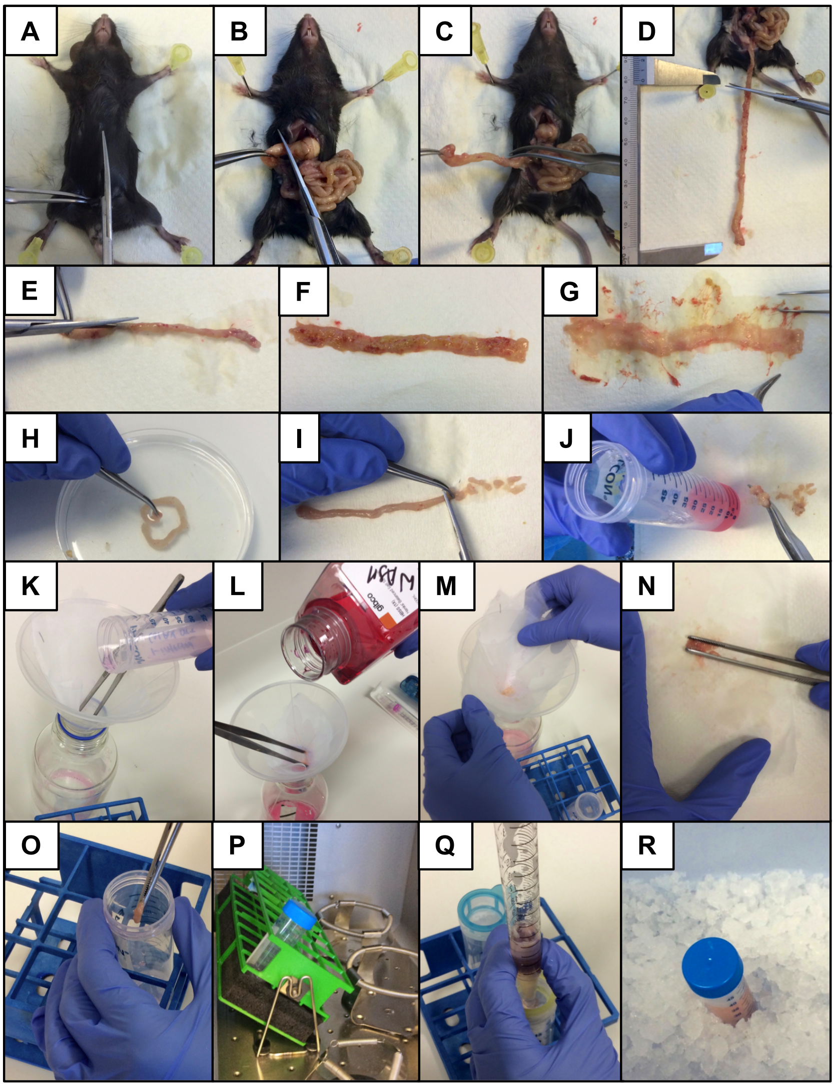
