## Supplementary Table 1 for "Single-cell analysis of intestinal immune cells during helminth infection"

**23-color spectral flow cytometry panel for the analysis of intestinal immune cells during helminth infection**

| | Channel | Emission | Marker | Fluorophore | Clone | Company | Catalog ID | optimal staining dilution | working conc. (µg/ml) |
|---|---|---|---|---|---|---|---|---|---|
| 1 | V1 | 421 | CD3 | BV421 | 145-2C11 | BD | 562600 | 1:100 | 2 |
| 2 | V3 | 450 | CD103 | eFluor450 | 2E7 | Thermofisher | 48-1031-82 | 1:100 | 2 |
| 3 | V4 | 480 | ki67 | BV480 | B56 | BD | 566109 | 1:200 | 0.01 |
| 4 | V5 | 510 | Ep-CAM(CD326) | BV510 | G8.8 | Biolegend | 118231 | 1:400 | 2 |
| 5 | V9 | 570 | Ly6C | BV570 | HK1.4 | Biolegend | 128030 | 1:200 | 2 |
| 6 | V10 | 605 | MHCII | BV605 | M5/114.15.2 | BD | 563413 | 1:400 | 2 |
| 7 | V11 | 650 | CD11c | BV650 | N418 | Biolegend | 117339 | 1:100 | 2 |
| 8 | V13 | 705 | DCAMKL1 | primary | polyclonal | Abcam | Ab31704 | 1:4000 | 10 |
| | | | | Qdot705 | | Thermofisher | Q-11461MP | 1:4000 | 10 |
| 9 | V14 | 750 | CD45 | BV750 | 30-F11 | Biolegend | 103157 | 1:1000 | 2 |
| 10 | V16 | 786 | Siglec-F | BV786 | E50-2440 | BD | 740956 | 1:200 | 2 |
| 11 | B2 | 520 | CD19 | AF488 | 6D5 | BioLegend | 115521 | 1:200 | 5 |
| 12 | B3 | 550 | FoxP3 | AF532 | FJK-16s | ThermoFisher | 58-5773-82 | 1:100 | 2 |
| 13 | B5 | 576 | Tbet | PE | 4B10 | BioLegend | 644809 | 1:100 | 2 |
| 14 | B6 | 610 | RORγt | PE-CF594 | Q31-378 | BD | 562684 | 1:100 | 2 |
| 15 | B8 | 668 | CD127 | PECy5 | A7R34 | BioLegend | 135016 | 1:100 | 2 |
| 16 | B9 | 680 | CD11b | PErCp Cy5.5 | M1/70 | BD | 550993 | 1:100 | 2 |
| 17 | B10 | 730 | CD4 | PerCP eFluor 710 | RM4-5 | ThermoFisher | 46-0042-82 | 1:1200 | 2 |
| 18 | B14 | 780 | CD64 | PE-Cy7 | X54-5/7.1 | BioLegend | 139314 | 1:100 | 2 |
| 19 | R2 | 660 | RELMa | APC | DS8RELM | ThermoFisher | 17-5441-82 | 1:100 | 2 |
| 20 | R2 | 668 | GATA3 | AF647 | L50-823 | BD | 560068 | 1:200 | 2 |
| 21 | R5 | 720 | Ly6G | AF700 | 1A8 | Biolegend | 127622 | 1:300 | 5 |
| 22 | R8 | 746 | ZOMBIE NIR | Live/death | - | Biolegend | 423106 | 1:1000 | |
| 23 | R8 | 780 | CD90.2 | APC-CY7 | 30-H12 | Biolegend | 105328 | 1:800 | 2 |

intracellular staining
