## Supplementary Figures 1-11 for "Single-cell analysis of intestinal immune cells during helminth infection"

| Digestion Protocol #1 | |
|---|---|
| 2x 15 min at 200 rpm | 1 mg/ml Collagenase VIII |
| HBSS/2 mM EDTA | 0.05 mg/ml DNAse |
| | 20 min at 200 rpm |

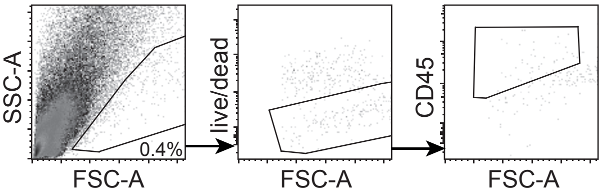


| Digestion Protocol #2 | |
|---|---|
| 4x 10 min at 200 rpm | 1 mg/ml Collagenase VIII |
| HBSS/2 mM EDTA | 0.05 mg/ml DNAse |
| | 20 min at 150 rpm |

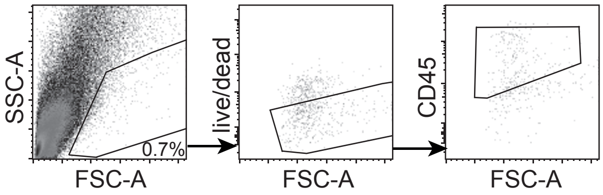


| Digestion Protocol #3 | |
|---|---|
| 4x 10 min at 200 rpm | 0.425 mg/ml Collagenase V |
| HBSS/2 mM EDTA | 0.625 mg/ml Collagenase D |
| | 1 mg/ml Dispase |
| | 0.05 mg/ml DNAse |
| | 15 min at 200 rpm |

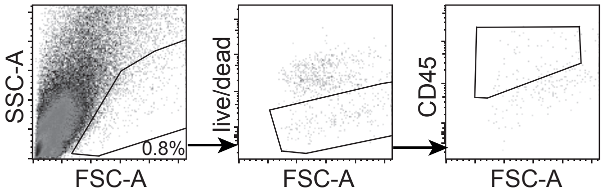


| Digestion Protocol #4 | |
|---|---|
| 3x 10 min at 200 rpm | 1 mg/ml Collagenase VIII |
| HBSS/2 mM EDTA | 0.425 mg/ml Collagenase V |
| | 0.625 mg/ml Collagenase D |
| | 20% FCS |
| | 0.05 mg/ml DNAse |
| | 10 min at 200 rpm |

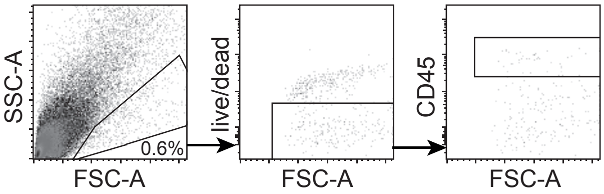


| Digestion Protocol #5 | |
|---|---|
| 4x 10 min at 200 rpm | 200 µg/ml Liberase TM |
| HBSS/2 mM EDTA | 20% FCS |
| | 0.05 mg/ml DNAse |
| | 20 min at 200 rpm |

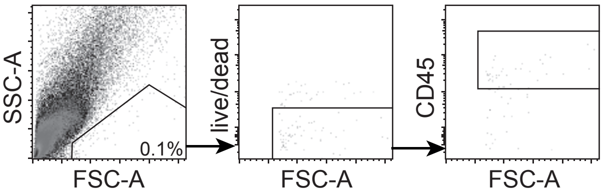


| Digestion Protocol #6 | |
|---|---|
| 4x 15 min at 200 rpm | 1 mg/ml Collagenase VIII |
| HBSS/2 mM EDTA | 20% FCS |
| | 0.05 mg/ml DNAse |
| | 25 min at 200 rpm |

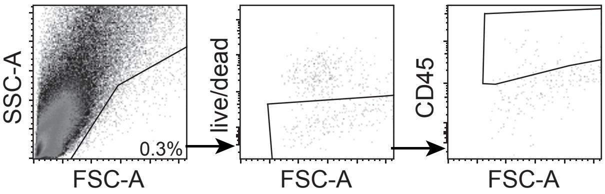


| Digestion Protocol #7 | |
|---|---|
| 2x 15 min at 200 rpm | 1 mg/ml Collagenase A |
| HBSS/2 mM EDTA | 20% FCS |
| | 0.05 mg/ml DNAse |
| | 20 min at 200 rpm |

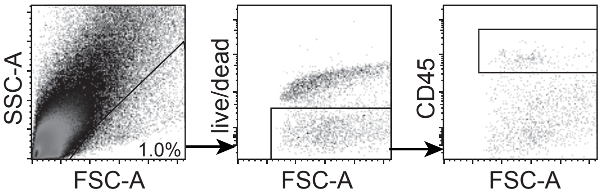


| Digestion Protocol #8 | |
|---|---|
| 4x 15 min at 200 rpm | 1 mg/ml Collagenase A |
| HBSS/2 mM EDTA | 20% FCS |
| | 0.05 mg/ml DNAse |
| | 25 min at 200 rpm |

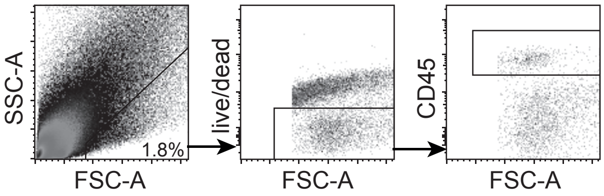

| | |
|---|---|
| <u>Digestion Protocol #9</u><br>6x 10 min at 200 rpm<br>HBSS/2 mM EDTA | 1 mg/ml Collagenase A<br>20% FCS<br>0.05 mg/ml DNAse<br>25 min at 200 rpm |

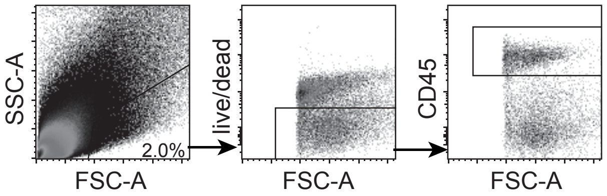


| | |
|---|---|
| <u>Digestion Protocol #10</u><br>2x 30 min at 200 rpm<br>HBSS/2 mM EDTA | 1 mg/ml Collagenase A<br>20% FCS<br>0.05 mg/ml DNAse<br>25 min at 200 rpm |

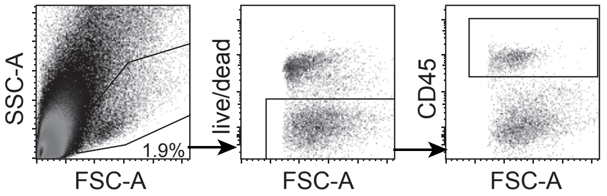


| | |
|---|---|
| <u>Digestion Protocol #11</u><br>1x 15 min at 200 rpm<br>HBSS/DTT<br>2x 15 min at 200 rpm<br>HBSS/2 mM EDTA | 1 mg/ml Collagenase A<br>20% FCS<br>0.05 mg/ml DNAse<br>25 min at 200 rpm |

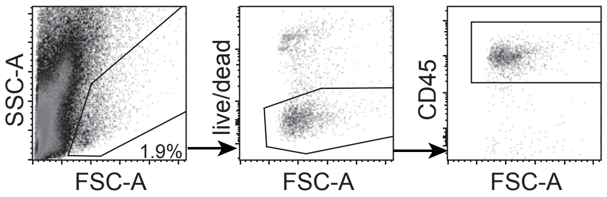


| | |
|---|---|
| <u>Digestion Protocol #12</u><br>3x 15 min at 200 rpm<br>HBSS/2 mM EDTA | 1 mg/ml Collagenase A<br>20% FCS<br>0.05 mg/ml DNAse<br>50 min at 200 rpm |

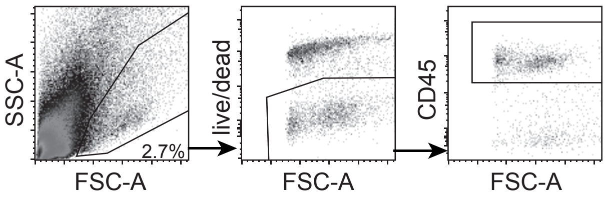


| | |
|---|---|
| <u>Digestion Protocol #13</u><br>3x 10 min at 200 rpm<br>HBSS/2 mM EDTA<br>+vortexing | 1 mg/ml Collagenase A<br>20% FCS<br>0.05 mg/ml DNAse<br>30 min at 200 rpm |

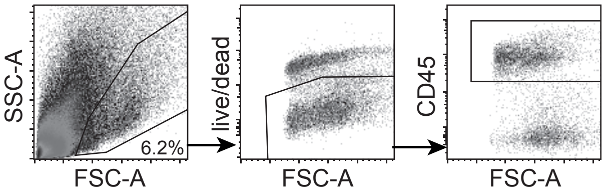


| | |
|---|---|
| <u>Digestion Protocol #14</u><br>3x 10 min at 200 rpm<br>HBSS/2 mM EDTA<br>+vortexing | 1 mg/ml Collagenase A<br>20% FCS<br>0.05 mg/ml DNAse<br>30 min at 200 rpm<br>40/80% Percoll gradient |

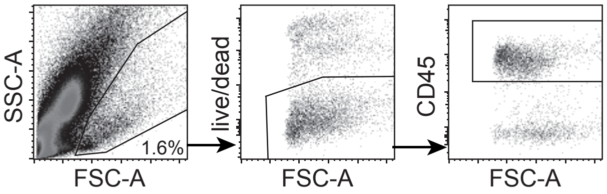

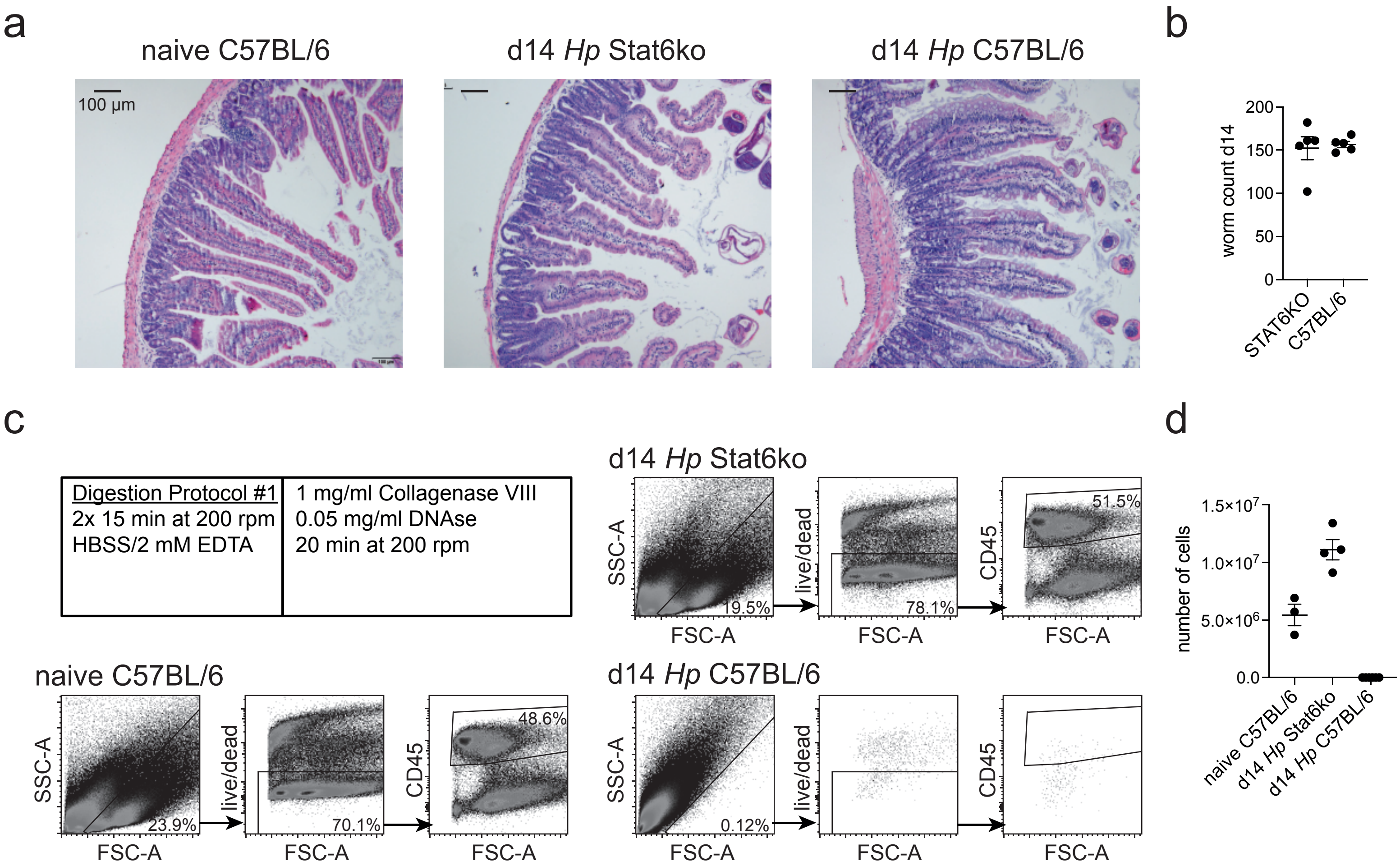

a
naive C57BL/6
d14 Hp Stat6ko
d14 Hp C57BL/6
100 µm
b
worm count d14
200
150
100
50
0
STAT6KO
C57BL/6
c
Digestion Protocol #1
2x 15 min at 200 rpm
HBSS/2 mM EDTA
1 mg/ml Collagenase VIII
0.05 mg/ml DNAse
20 min at 200 rpm
d14 Hp Stat6ko
SSC-A
FSC-A
19.5%
live/dead
78.1%
CD45
51.5%
naive C57BL/6
23.9%
70.1%
48.6%
d14 Hp C57BL/6
0.12%
d
number of cells
1.5×10^7
1.0×10^7
5.0×10^6
0.0
naive C57BL/6
d14 Hp Stat6ko
d14 Hp C57BL/6

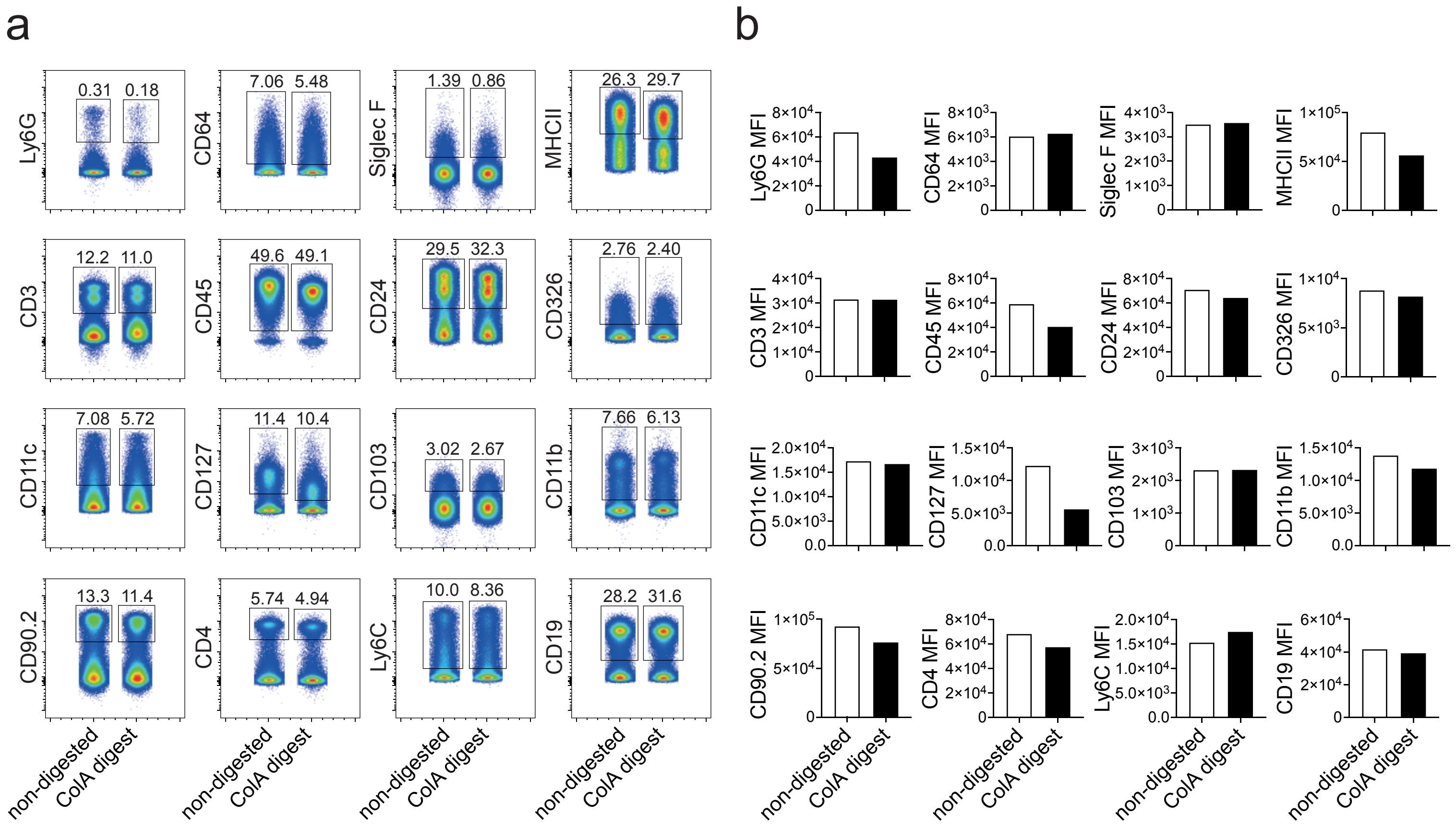
a
b
Ly6G
0.31
0.18
CD64
7.06
5.48
Siglec F
1.39
0.86
MHCII
26.3
29.7
CD3
12.2
11.0
CD45
49.6
49.1
CD24
29.5
32.3
CD326
2.76
2.40
CD11c
7.08
5.72
CD127
11.4
10.4
CD103
3.02
2.67
CD11b
7.66
6.13
CD90.2
13.3
11.4
CD4
5.74
4.94
Ly6C
10.0
8.36
CD19
28.2
31.6
non-digested
ColA digest
Ly6G MFI
CD64 MFI
Siglec F MFI
MHCII MFI
CD3 MFI
CD45 MFI
CD24 MFI
CD326 MFI
CD11c MFI
CD127 MFI
CD103 MFI
CD11b MFI
CD90.2 MFI
CD4 MFI
Ly6C MFI
CD19 MFI

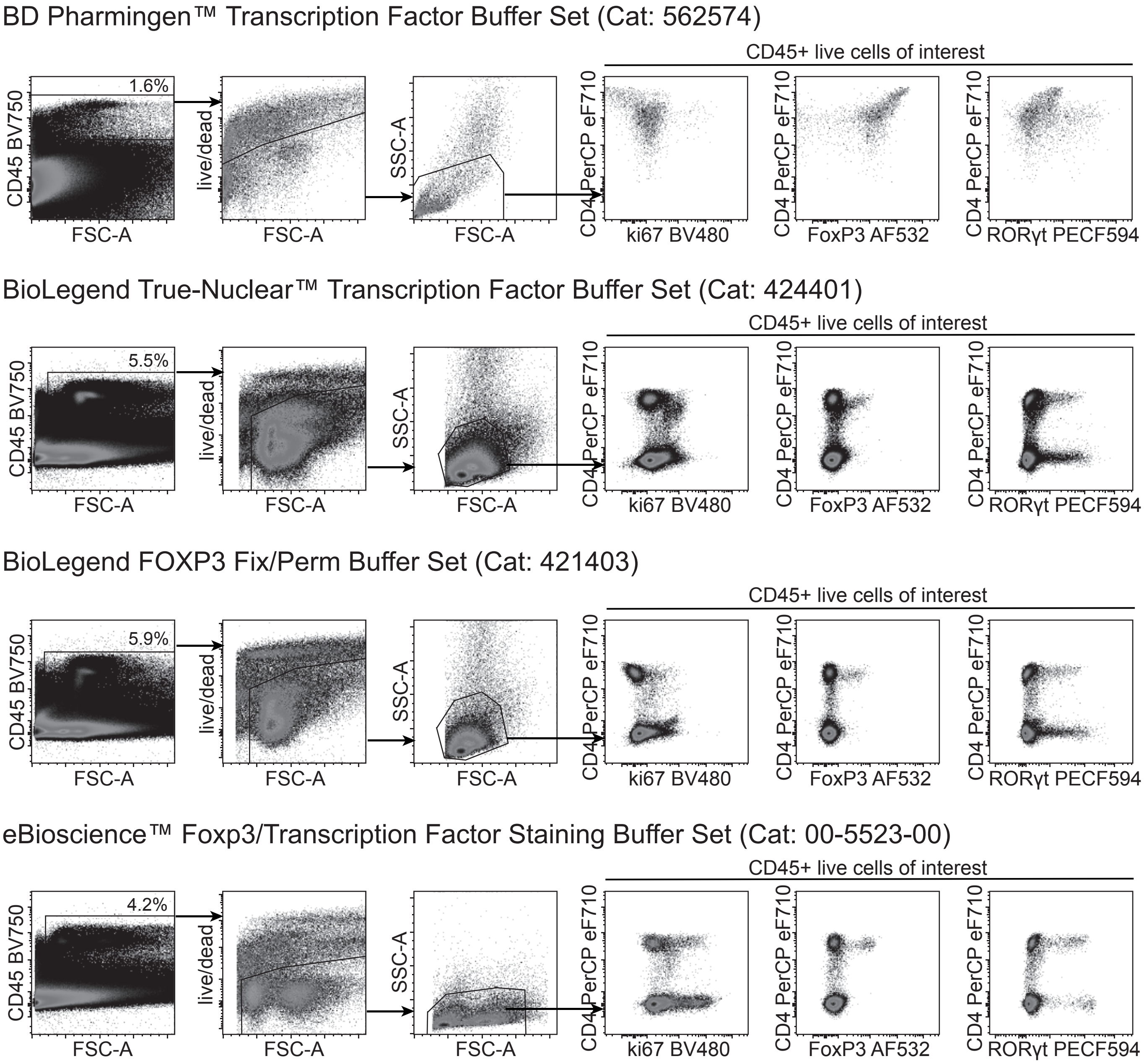
BD Pharmingen™ Transcription Factor Buffer Set (Cat: 562574)
CD45+ live cells of interest
1.6%
CD45 BV750
FSC-A
live/dead
FSC-A
SSC-A
FSC-A
CD4 PerCP eF710
ki67 BV480
CD4 PerCP eF710
FoxP3 AF532
CD4 PerCP eF710
RORγt PECF594
BioLegend True-Nuclear™ Transcription Factor Buffer Set (Cat: 424401)
CD45+ live cells of interest
5.5%
CD45 BV750
FSC-A
live/dead
FSC-A
SSC-A
FSC-A
CD4 PerCP eF710
ki67 BV480
CD4 PerCP eF710
FoxP3 AF532
CD4 PerCP eF710
RORγt PECF594
BioLegend FOXP3 Fix/Perm Buffer Set (Cat: 421403)
CD45+ live cells of interest
5.9%
CD45 BV750
FSC-A
live/dead
FSC-A
SSC-A
FSC-A
CD4 PerCP eF710
ki67 BV480
CD4 PerCP eF710
FoxP3 AF532
CD4 PerCP eF710
RORγt PECF594
eBioscience™ Foxp3/Transcription Factor Staining Buffer Set (Cat: 00-5523-00)
CD45+ live cells of interest
4.2%
CD45 BV750
FSC-A
live/dead
FSC-A
SSC-A
FSC-A
CD4 PerCP eF710
ki67 BV480
CD4 PerCP eF710
FoxP3 AF532
CD4 PerCP eF710
RORγt PECF594

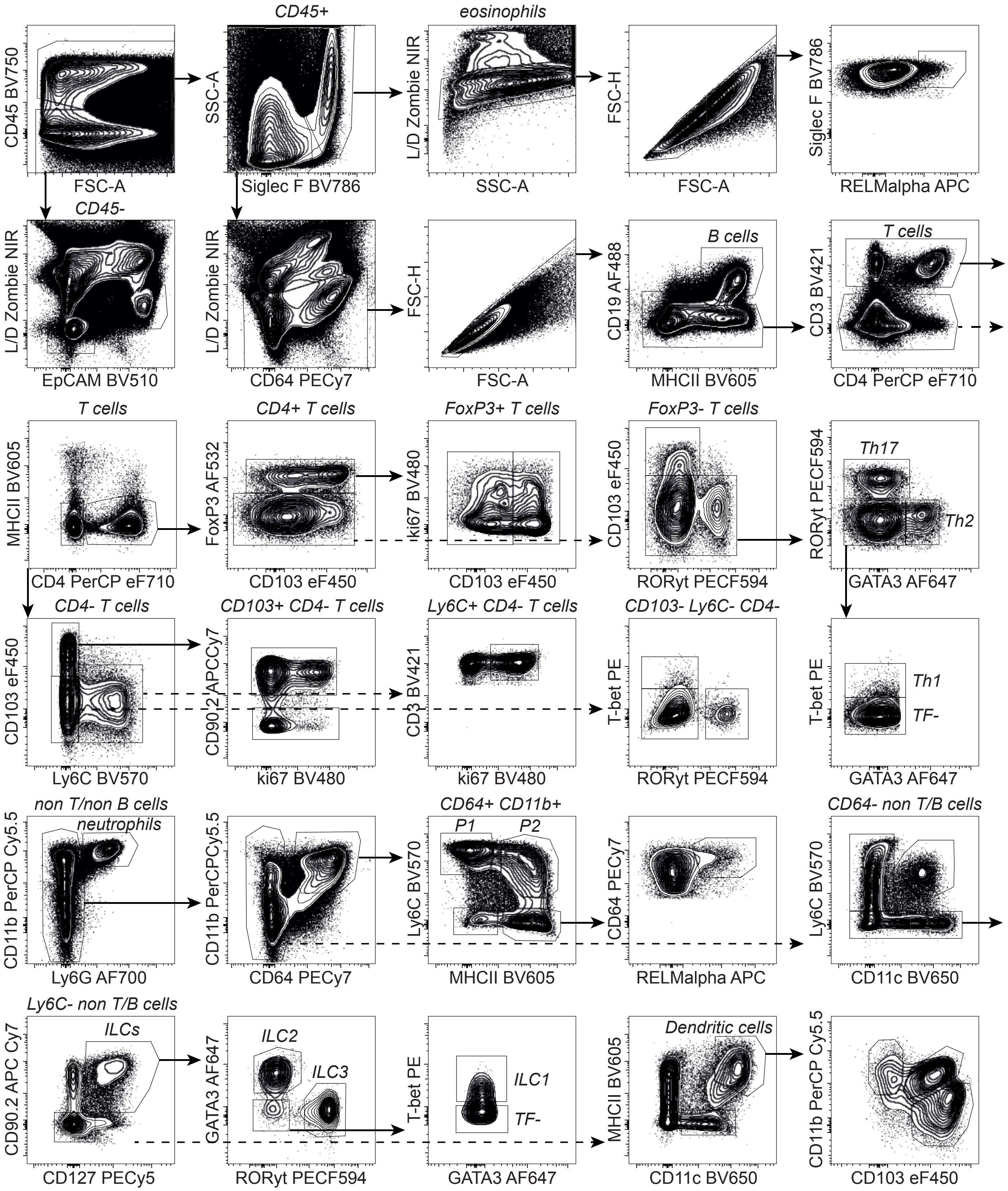
CD45 BV750
FSC-A
CD45+
SSC-A
Siglec F BV786
eosinophils
L/D Zombie NIR
SSC-A
FSC-H
FSC-A
Siglec F BV786
RELMalpha APC
CD45-
L/D Zombie NIR
EpCAM BV510
L/D Zombie NIR
CD64 PECy7
FSC-H
FSC-A
B cells
CD19 AF488
MHCII BV605
T cells
CD3 BV421
CD4 PerCP eF710
T cells
MHCII BV605
CD4 PerCP eF710
CD4+ T cells
FoxP3 AF532
CD103 eF450
FoxP3+ T cells
ki67 BV480
CD103 eF450
FoxP3- T cells
CD103 eF450
RORyt PECF594
Th17
RORyt PECF594
Th2
GATA3 AF647
CD4- T cells
CD103 eF450
Ly6C BV570
CD103+ CD4- T cells
CD90.2 APCCy7
ki67 BV480
Ly6C+ CD4- T cells
CD3 BV421
ki67 BV480
CD103- Ly6C- CD4-
T-bet PE
RORyt PECF594
T-bet PE
Th1
TF-
GATA3 AF647
non T/non B cells
neutrophils
CD11b PerCP Cy5.5
Ly6G AF700
CD11b PerCPCy5.5
CD64 PECy7
CD64+ CD11b+
P1
P2
Ly6C BV570
MHCII BV605
CD64 PECy7
RELMalpha APC
CD64- non T/B cells
Ly6C BV570
CD11c BV650
Ly6C- non T/B cells
ILCs
CD90.2 APC Cy7
CD127 PECy5
ILC2
ILC3
GATA3 AF647
RORyt PECF594
T-bet PE
ILC1
TF-
GATA3 AF647
Dendritic cells
MHCII BV605
CD11c BV650
CD11b PerCP Cy5.5
CD103 eF450

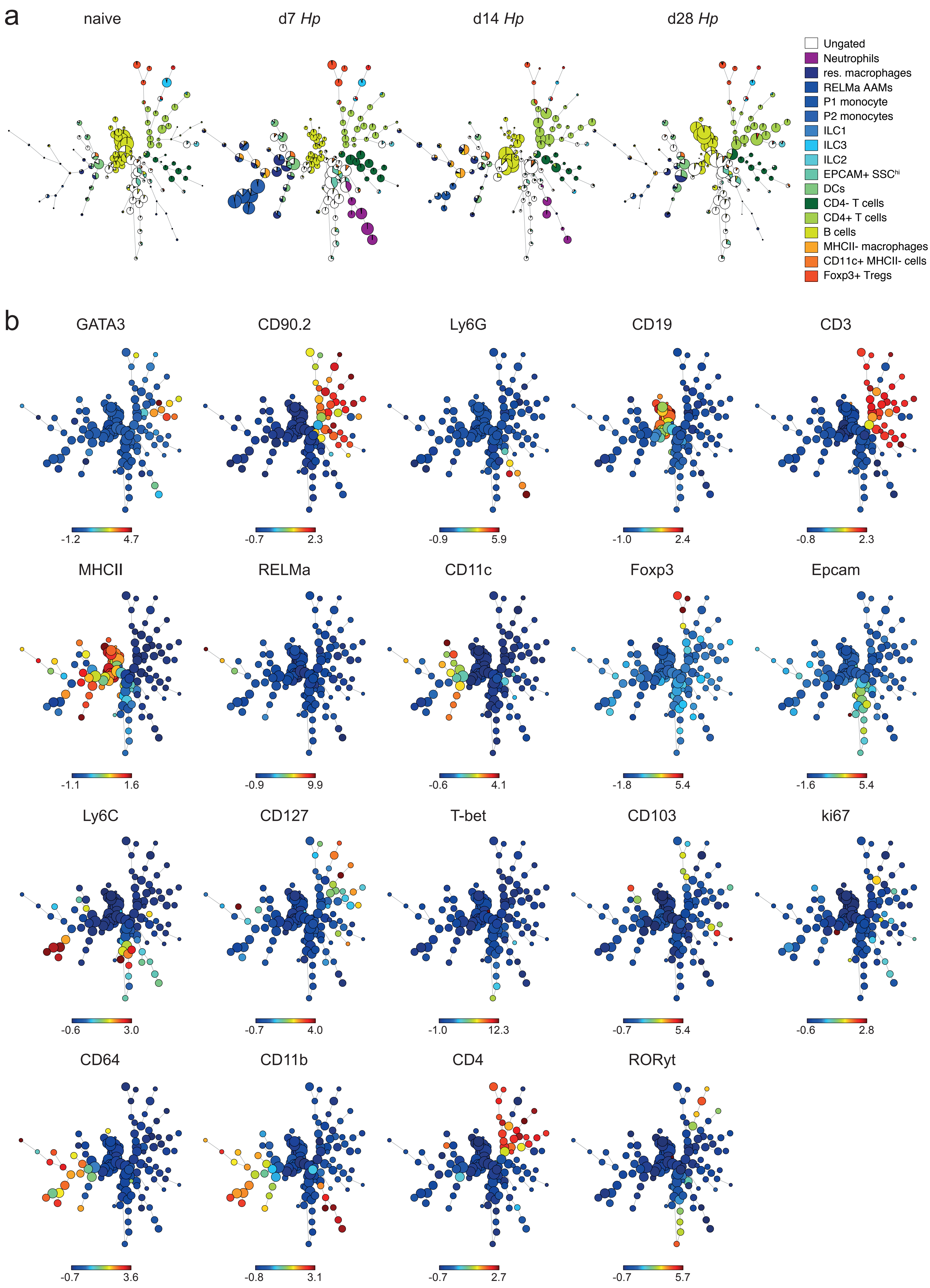
a
naive
d7 Hp
d14 Hp
d28 Hp
Ungated
Neutrophils
res. macrophages
RELMa AAMs
P1 monocyte
P2 monocytes
ILC1
ILC3
ILC2
EPCAM+ SSChi
DCs
CD4- T cells
CD4+ T cells
B cells
MHCII- macrophages
CD11c+ MHCII- cells
Foxp3+ Tregs
b
GATA3
-1.2 4.7
CD90.2
-0.7 2.3
Ly6G
-0.9 5.9
CD19
-1.0 2.4
CD3
-0.8 2.3
MHCII
-1.1 1.6
RELMa
-0.9 9.9
CD11c
-0.6 4.1
Foxp3
-1.8 5.4
Epcam
-1.6 5.4
Ly6C
-0.6 3.0
CD127
-0.7 4.0
T-bet
-1.0 12.3
CD103
-0.7 5.4
ki67
-0.6 2.8
CD64
-0.7 3.6
CD11b
-0.8 3.1
CD4
-0.7 2.7
RORyt
-0.7 5.7

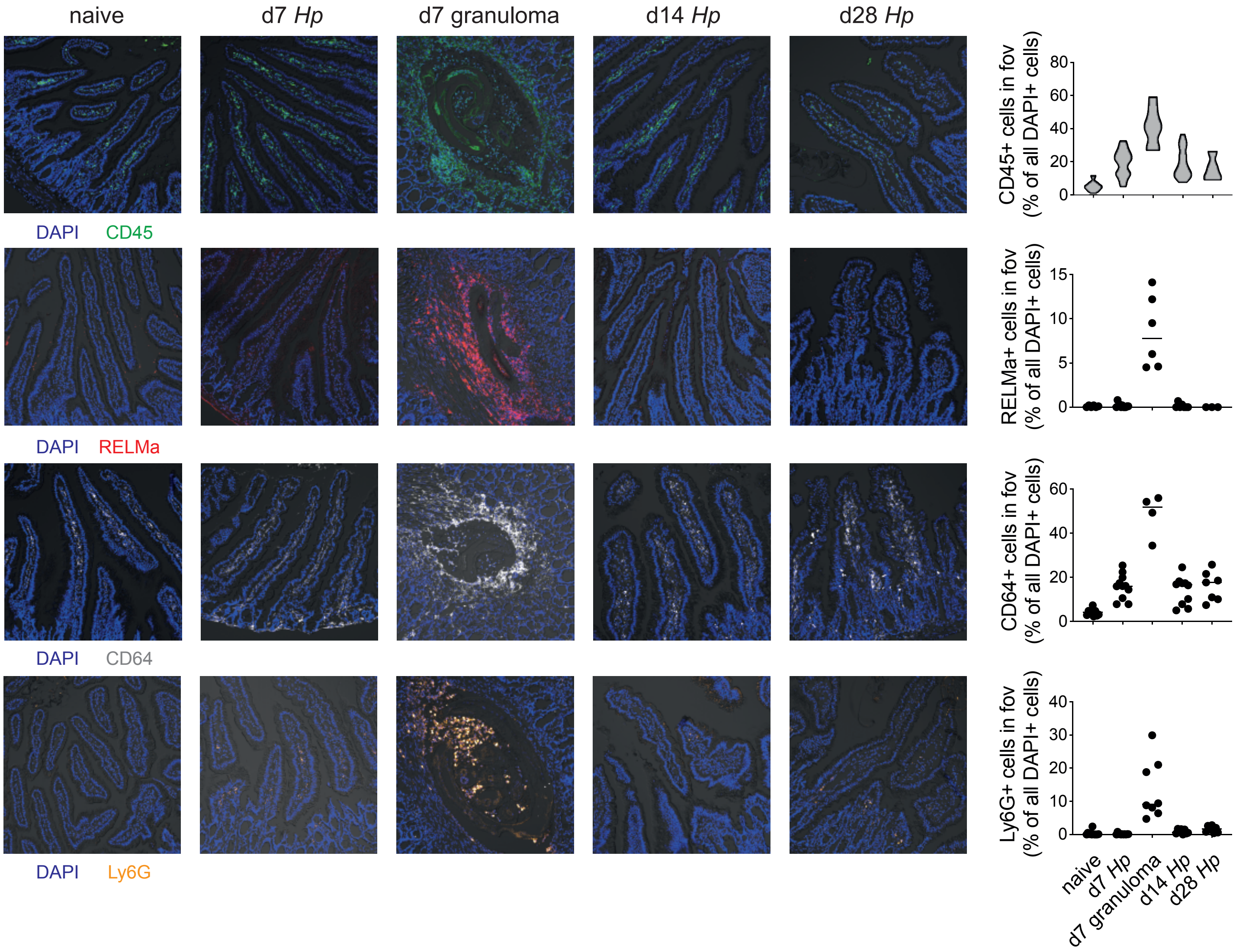
naive
d7 Hp
d7 granuloma
d14 Hp
d28 Hp
DAPI CD45
DAPI RELMa
DAPI CD64
DAPI Ly6G
CD45+ cells in fov (% of all DAPI+ cells)
RELMa+ cells in fov (% of all DAPI+ cells)
CD64+ cells in fov (% of all DAPI+ cells)
Ly6G+ cells in fov (% of all DAPI+ cells)
0
20
40
60
80
0
5
10
15
0
20
40
60
0
10
20
30
40
naive
d7 Hp
d7 granuloma
d14 Hp
d28 Hp

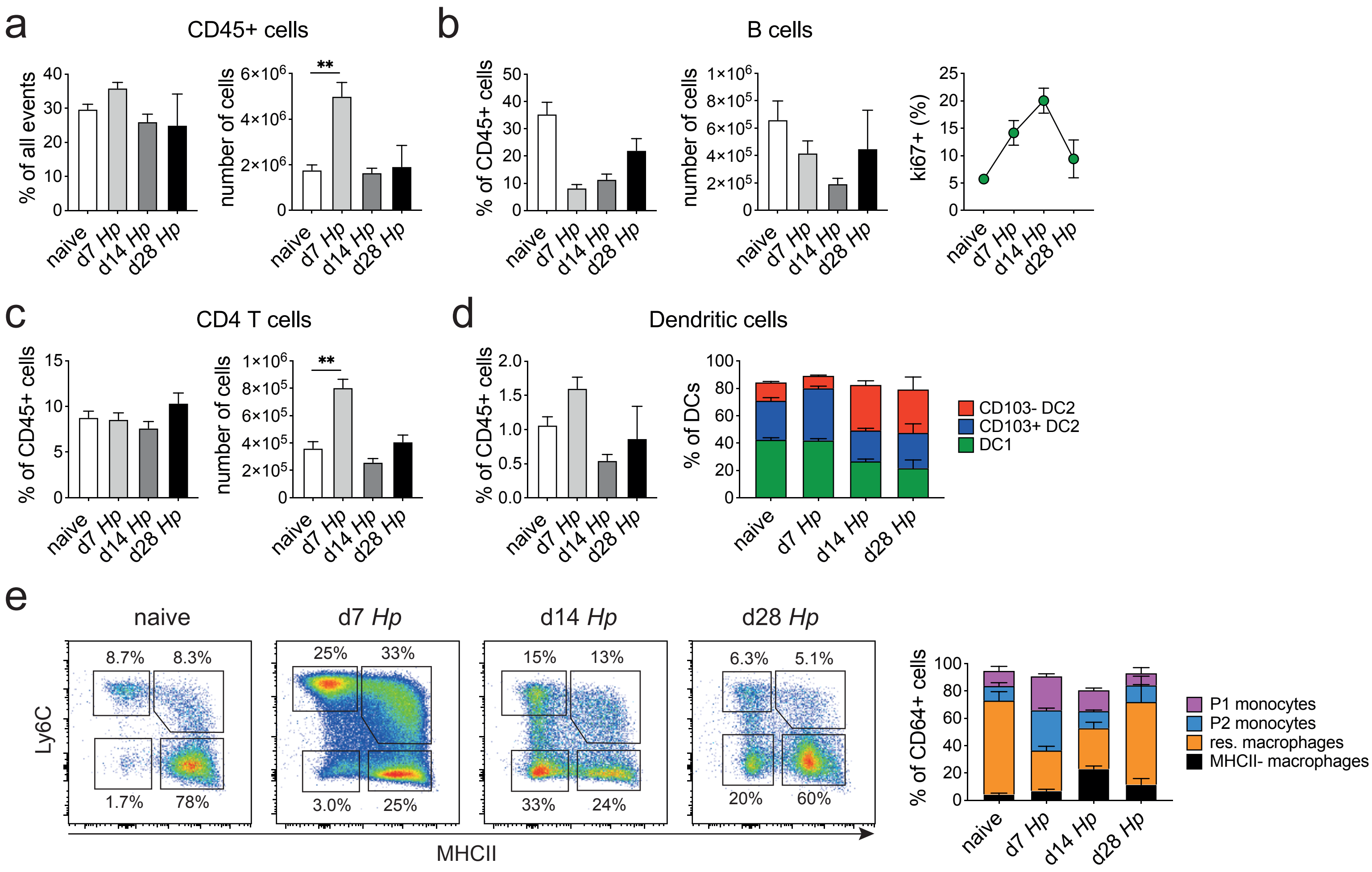
a
CD45+ cells
% of all events
number of cells
**
b
B cells
% of CD45+ cells
number of cells
ki67+ (%)
c
CD4 T cells
% of CD45+ cells
number of cells
**
d
Dendritic cells
% of CD45+ cells
% of DCs
CD103- DC2
CD103+ DC2
DC1
e
naive
d7 Hp
d14 Hp
d28 Hp
Ly6C
MHCII
8.7%
8.3%
1.7%
78%
25%
33%
3.0%
25%
15%
13%
33%
24%
6.3%
5.1%
20%
60%
% of CD64+ cells
P1 monocytes
P2 monocytes
res. macrophages
MHCII- macrophages
naive
d7 Hp
d14 Hp
d28 Hp

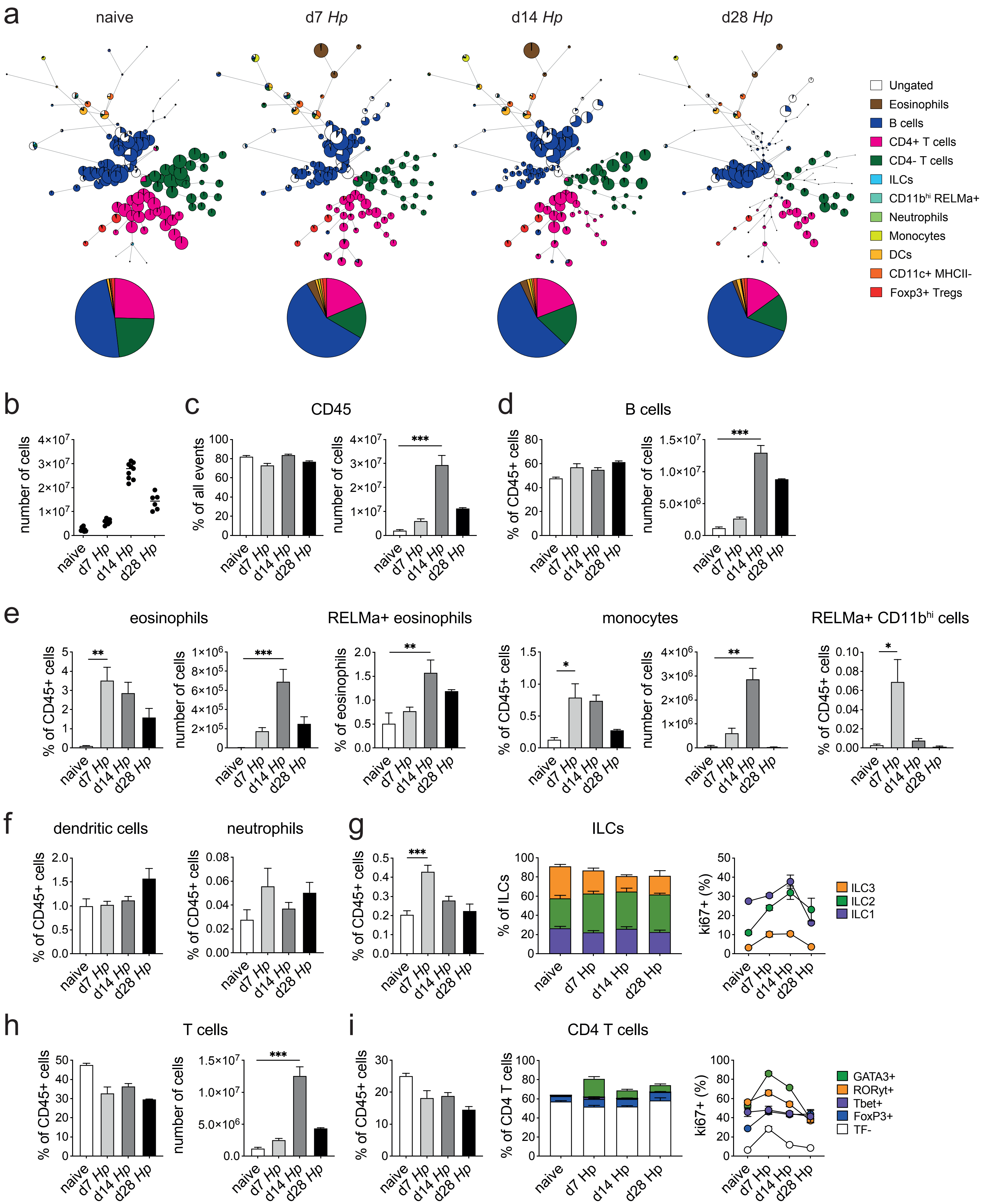
a
naive
d7 Hp
d14 Hp
d28 Hp
Ungated
Eosinophils
B cells
CD4+ T cells
CD4- T cells
ILCs
CD11b^hi RELMa+
Neutrophils
Monocytes
DCs
CD11c+ MHCII-
Foxp3+ Tregs
b
number of cells
c
CD45
% of all events
number of cells
d
B cells
% of CD45+ cells
number of cells
e
eosinophils
RELMa+ eosinophils
monocytes
RELMa+ CD11b^hi cells
% of CD45+ cells
number of cells
% of eosinophils
f
dendritic cells
neutrophils
g
ILCs
% of ILCs
ki67+ (%)
ILC3
ILC2
ILC1
h
T cells
i
CD4 T cells
% of CD4 T cells
GATA3+
RORyt+
Tbet+
FoxP3+
TF-
naive
d7 Hp
d14 Hp
d28 Hp

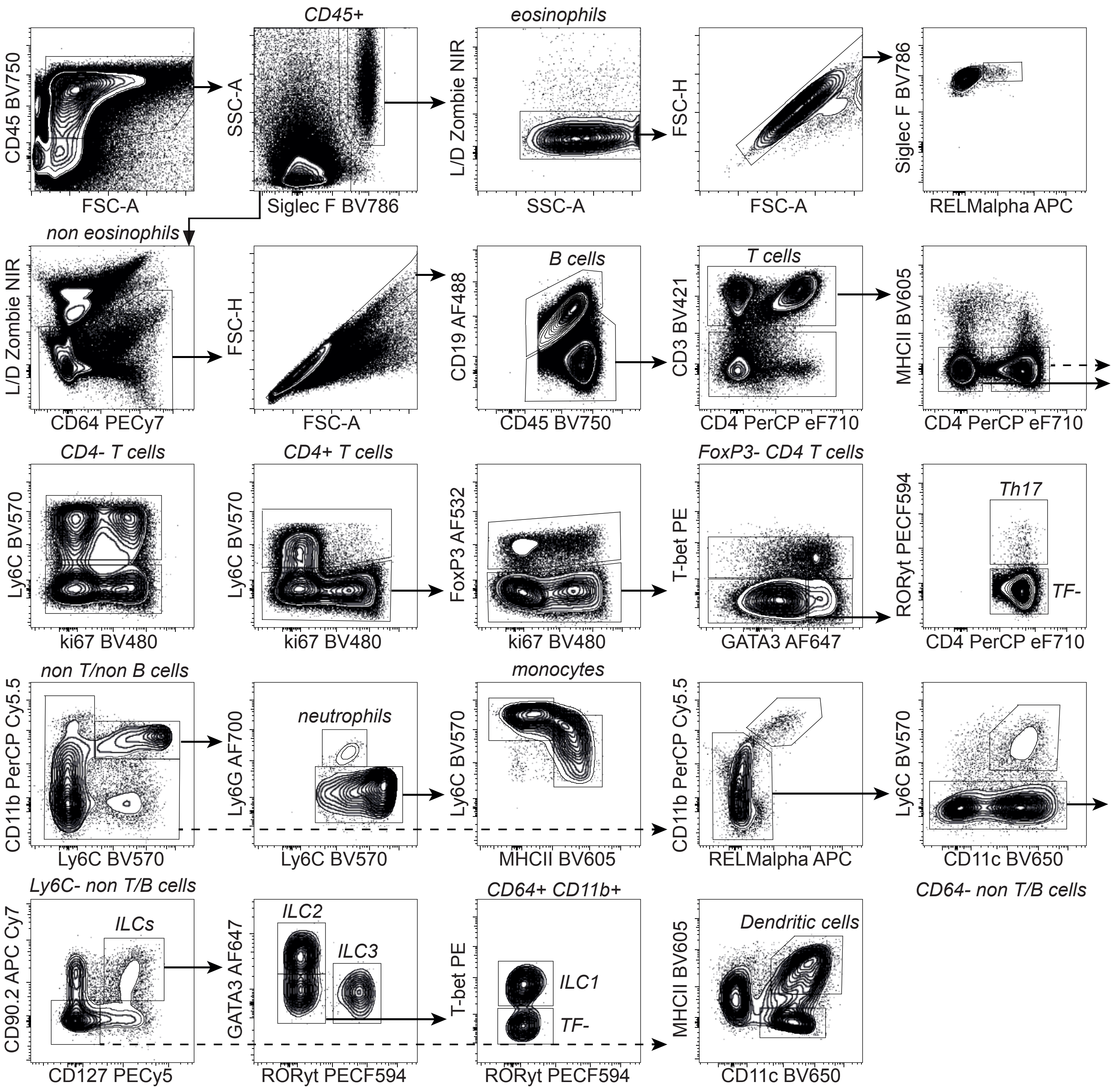
CD45 BV750
FSC-A
CD45+
SSC-A
Siglec F BV786
eosinophils
L/D Zombie NIR
SSC-A
FSC-H
FSC-A
Siglec F BV786
RELMalpha APC
non eosinophils
L/D Zombie NIR
CD64 PECy7
FSC-H
FSC-A
B cells
CD19 AF488
CD45 BV750
T cells
CD3 BV421
CD4 PerCP eF710
MHCII BV605
CD4 PerCP eF710
CD4- T cells
Ly6C BV570
ki67 BV480
CD4+ T cells
Ly6C BV570
ki67 BV480
FoxP3 AF532
ki67 BV480
FoxP3- CD4 T cells
T-bet PE
GATA3 AF647
Th17
RORyt PECF594
TF-
CD4 PerCP eF710
non T/non B cells
CD11b PerCP Cy5.5
Ly6C BV570
neutrophils
Ly6G AF700
Ly6C BV570
monocytes
Ly6C BV570
MHCII BV605
CD11b PerCP Cy5.5
RELMalpha APC
Ly6C BV570
CD11c BV650
CD64- non T/B cells
Ly6C- non T/B cells
ILCs
CD90.2 APC Cy7
CD127 PECy5
ILC2
ILC3
GATA3 AF647
RORyt PECF594
CD64+ CD11b+
ILC1
TF-
T-bet PE
RORyt PECF594
Dendritic cells
MHCII BV605
CD11c BV650
